# Mass spectrometric based detection of protein nucleotidylation in the RNA polymerase of SARS-CoV-2

**DOI:** 10.1101/2020.10.07.330324

**Authors:** Brian J. Conti, Robert N. Kirchdoerfer, Michael R. Sussman

## Abstract

Coronaviruses, like SARS-CoV-2, encode a nucleotidyl transferase in the N-terminal NiRAN domain of the <u>n</u>on-<u>s</u>tructural <u>p</u>rotein (nsp) 12 protein within the <u>R</u>NA dependent <u>R</u>NA <u>p</u>olymerase (RdRP) ^1-3^. Though the substrate targets of the viral nucleotidyl transferase are unknown, NiRAN active sites are highly conserved and essential for viral replication ^3^. We show, for the first time, the detection and sequence location of GMP-modified amino acids in nidovirus RdRP-associated proteins using heavy isotope-assisted MS and MS/MS peptide sequencing. We identified lys-143 in the equine arteritis virus (EAV) protein, nsp7, as a primary site of nucleotidylation in vitro that uses a phosphoramide bond to covalently attach with GMP. In SARS-CoV-2 replicase proteins, we demonstrate a unique O-linked GMP attachment on nsp7 ser-1, whose formation required the presence of nsp12. It is clear that additional nucleotidylation sites remain undiscovered, which includes the possibility that nsp12 itself may form a transient GMP adduct in the NiRAN active site that has eluted detection in these initial studies due to instability of the covalent attachment. Our results demonstrate new strategies for detecting GMP-peptide linkages that can be adapted for higher throughput screening using mass spectrometric technologies. These data are expected to be important for a rapid and timely characterization of a new enzymatic activity in SARS-CoV-2 that may be an attractive drug target aimed at limiting viral replication in infected patients.

---

The world is faced with a pandemic disease, COVID-19, caused by the emergence and global spread of a new species of coronavirus. Although it is clear that lethality of the disease progression is often caused by pulmonary problems associated with severe respiratory symptoms, this virus continues to surprise the medical community with new pathologies that are creating challenges for effective treatment. While the causative agent of COVID-19, SARS-CoV-2, is similar to other coronaviruses that infect humans and animals, many molecular details of this virus’s macromolecular structures and functions remain unknown. Immunological approaches to neutralize the virus, using both passive (e.g. injection of antibodies into patients) or active (e.g., injection of DNA, mRNA or protein that generates a neutralizing immune response) immunity are being pursued by scientists in both the private and public sectors. Since it is unclear whether any of these methods will succeed, it is prudent to explore other approaches to treat this disease, including drug therapy. Small molecule drug therapy has been highly successful in controlling HIV-1 infections and curing infections of hepatitis C virus ^4-7^. Creating small molecule drugs that target the SARS-CoV-2 replication machinery would provide a complementary approach to ongoing vaccine development efforts as well as preparing the world for future outbreaks caused by emerging coronaviruses.

A logical target for drugs that inhibit the virus is the conserved protein machinery responsible for replication of the viral RNA genome. Targeting proteins associated with the viral RNA-dependent RNA Polymerase (RdRP) is particularly attractive given the absence of similar RNA replication machinery in human cells. The small molecule Remsedevir targets the coronavirus replication machinery causing premature RNA chain termination ^8^. Remdesivir is now undergoing clinical trials and has been approved for compassionate use in the treatment of COVID-19. Discovery of new enzymatic and protein binding activities in SARS-CoV-2 is a critical part in elucidating the viral life cycle and importantly, in developing novel strategies to combat this disease as well as potentially new emerging viruses that use similar enzymes.

Here, we characterize the nucleotidyl transferase activity in SARS-CoV-2 replicase proteins by the detection of GMP-protein adducts using mass spectrometric technologies. The SARS-CoV-2 RdRP nsp12 mediated the nucleotidylation of nsp7 and nsp8, which bind near the nsp12 RNA polymerase active site and act as essential co-factors in RNA synthesis ^2,9^. Similar nucleotidylation activity was recently discovered in the RdRP of equine arteritis virus (EAV), nsp9, within the newly defined NiRAN domain (Nidovirus RdRP associated nucleotidyltransferase) that is also conserved in SARS-CoV-2 ^1-3^. Mutation of residues that were critical in EAV nsp9 nucleotidyl transferase activity, as well as mutation of the corresponding residues in SARS-CoV nsp12, eliminated viral propagation ^3^. However, there has been no definitive direct chemical analysis of the bond covalently linking the nucleotides to the protein, nor the exact chemical structure of the modification and the amino acids in which it occurs.

In order to utilize mass spectrometric technologies to provide definitive chemical data on the nucleotidylation event in nidoviruses, we first confirmed the earlier report of nucleotidyl transferase activity in EAV. We incubated nsp9 (Mr∼76,800) together with α-^32^P-GTP and a 2-4 molar excess of nsp7 (Mr∼25,200), which binds nsp9 and plays an important role in viral propagation^10,11^. Nucleotidyated products were analyzed by SDS-PAGE followed by Coomassie staining and autoradiography (Fig. 1). The same analysis was performed on SARS-CoV-2 nsp7 (Mr∼9,300), nsp8 (Mr∼21,900) and nsp12 (Mr∼106,700) that represent the minimal core protein subunits comprising the RdRP (note that by convention, the numbering of nonstructural proteins that comprise the RdRP of EAV is independent of that in SARS-CoV-2 and reflects their position within the viral polyprotein, not their function). The proteins were kept at low temperatures and neutral pH to stabilize labile, phosphoramide linkages between GMP and protein (Extended Data Fig. 1) ^3,12,13^. Although EAV nsp9 alone was sufficient to observe protein nucleotidylation^3^, the addition of nsp7 resulted in higher levels of radionucleotide incorporation into both proteins (Fig. 1, Extended Data Fig. 2). In contrast, full length SARS-CoV-2 nsp12 never incorporated radioactivity itself but was necessary to observe nucleotidylation on nsp7 and nsp8. In EAV nsp9, residue K94 is critical for nucleotidyl transferase activity. Mutation of the homologous K73 residue in SARS-CoV, which is also conserved in SARS-CoV-2, abolishes viral propagation in Vero-E6 cells ^3,14^. In our assay, a mutation of SARS-CoV-2 nsp12 that replaced a basic amino acid with a neutral amino acid, i.e. K73A, abolished all incorporation of radioactivity in nsp7 and nsp8, further demonstrating the central role of this nsp12 RdRP protein in nucleotidyation. Promiscuous labeling of a non-relevant protein, bovine serum albumin (BSA), was not observed suggesting nsp12-mediated nucleotidylation exhibits substrate specificity, possibly through close localization of proteins within the macromolecular polymerase complex.

**Figure 1.**
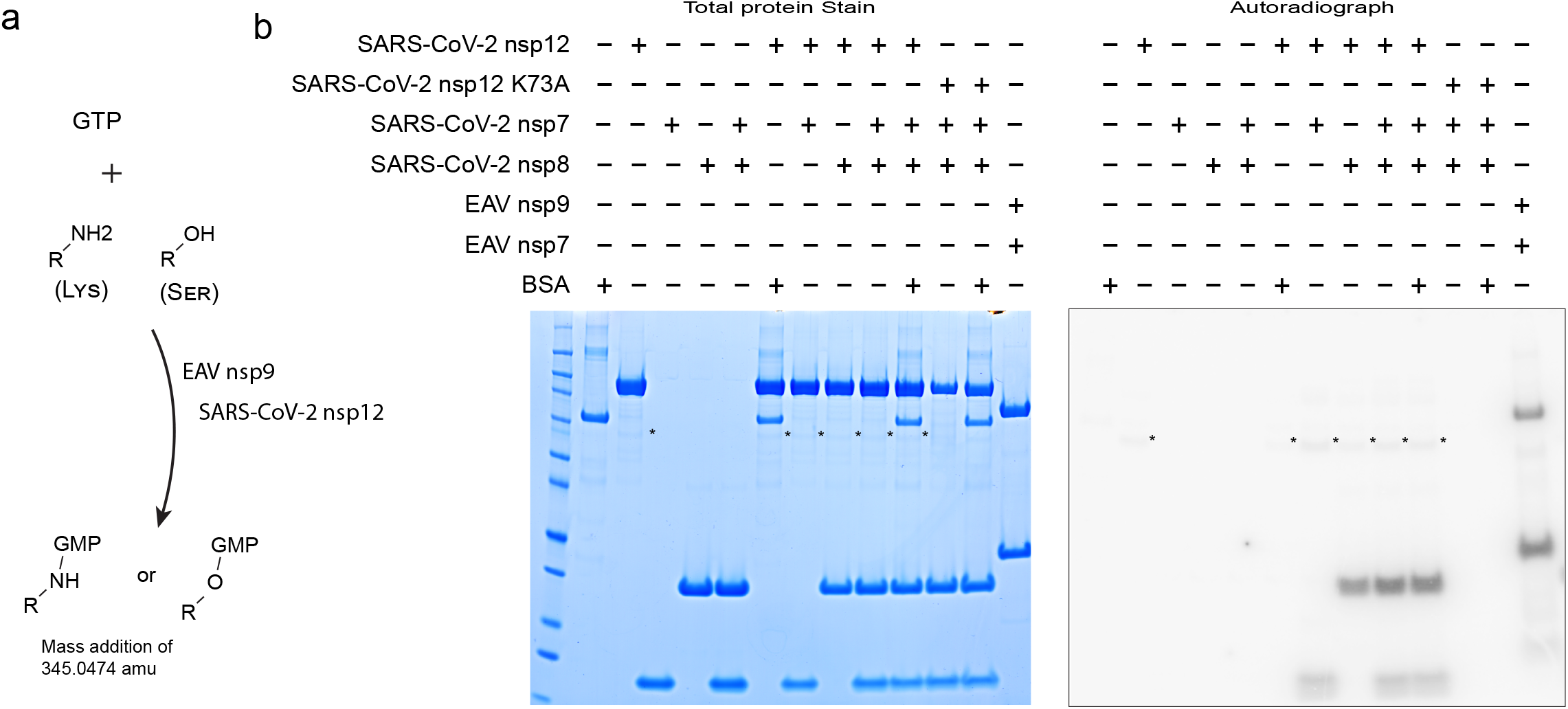
SARS-CoV-2 and EAV RdRP proteins are radiolabeled by incubation with α-^32^P-GTP. a, Summary of general findings observed in this study. b, SDS-PAGE analysis of nucleotidylation reactions that contained EAV nsp7 and RdRP nsp9, or alternatively the SARS-CoV-2 nsp7, nsp8, and/or RdRP nsp12 combinations along with BSA as indicated. Gels were stained with Coomassie to visualize total protein (left panel) or subjected to autoradiography to visualize proteins that were radiolabeled after incubation with α-^32^P-GTP (right panel). Asterisks are shown to indicate location of a faint band observed in both the Coomassie stained gel and autoradiograph.

We next utilized heavy isotope incorporation and the high mass accuracy of a tribrid Orbitrap based mass spectrometer to confirm that the transfer of radioactivity to the nsp proteins was a result of nucleotidylation. We performed nucleotidylation reactions in the absence of nucleotide, in the presence of natural GTP or, alternatively, GTP containing the heavy isotopes ^15^N or ^13^C. This allowed us to identify GMP-labeled peptide peaks using LC-MS alone, independent of MS/MS spectrum acquisition and matching. Unlabeled nsp peptides that were present in all sample injections ran at similar retention times (Fig. 2a, Extended Data Fig. 3). LC-MS peaks that corresponded to GMP-labeled peptides were identified by two criteria (Fig. 2b): (1) their absence in unlabeled sample injections, and (2) the presence of appropriately mass-shifted peptides at the same retention time in samples that were labeled with ^15^N- and ^13^C-GTP. Although a typical SARS-CoV-2 sample had ∼13,000 different LC-MS peaks, only three peaks were found that met the above criteria of GMP-labeled peptides (Extended Data Fig. 4, Supplementary Tables 1 and 2). Similarly, the sample set for EAV nsp proteins contained ∼91,000 peaks, but only 50 peaks satisfied the specified criteria as GMP modified candidates (Extended Data Fig.5, Supplementary Table 3).

**Figure 2.**
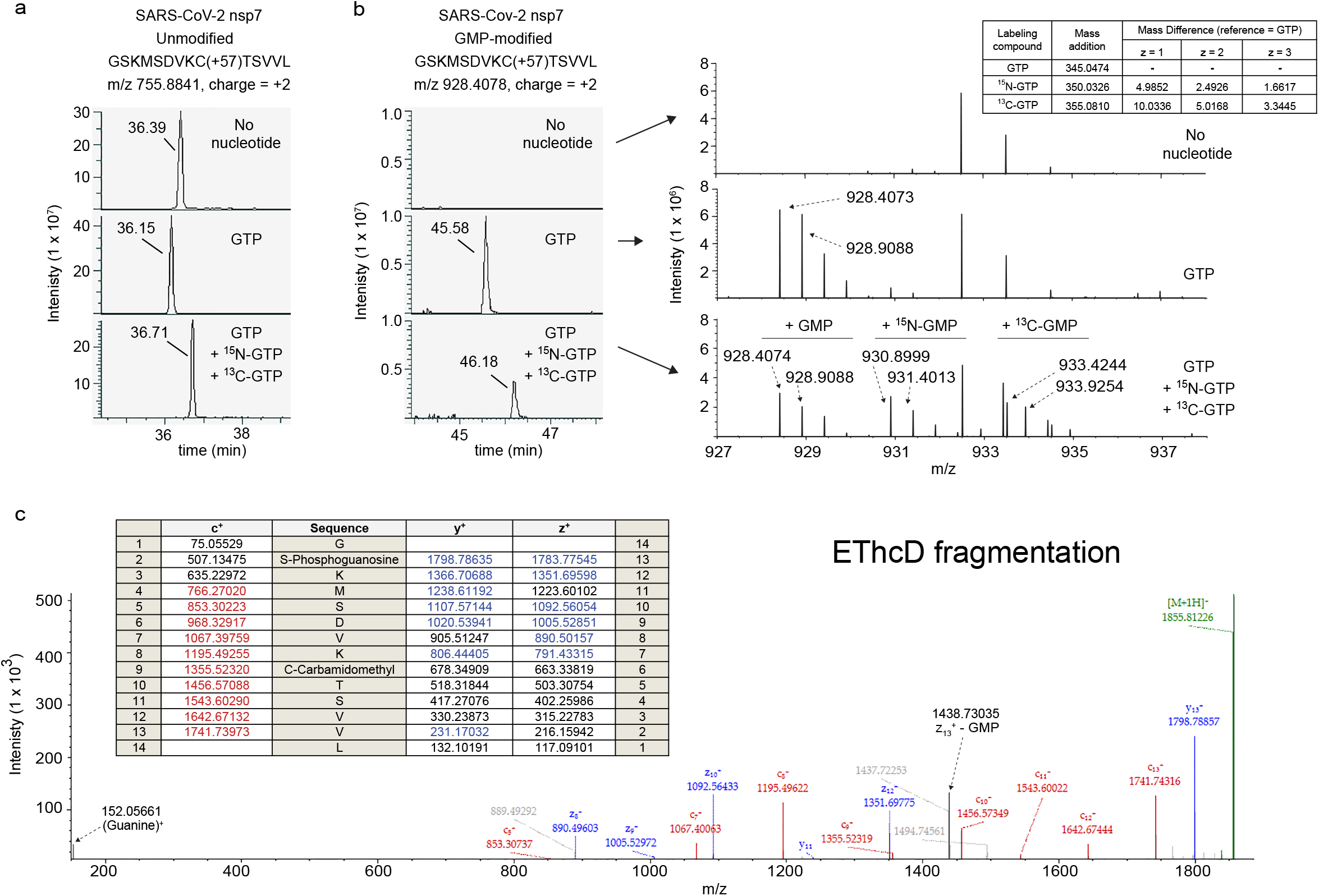
Detection of a GMP-modified SARS-CoV-2 peptide using LC-MS/MS. a, Extracted Ion chromatograms at the indicated m/z (+/- 5 ppm) show the precursor peptide MS1 signals versus retention time for the SARS-CoV-2 nsp7 peptide 1-14 with a charge of +2. Signals are shown for samples that were incubated without GTP (no nucleotide), with GTP, or with a mixture of GTP, ^15^N-GTP and ^13^C-GTP (top, middle and lower panels, respectively). b, MS1 signals at the m/z of 928.4078 for the same samples are shown (left panel). This mass corresponds to the GMP-modified version of the same peptide with a charge state of +2. The average mass spectrum across these peaks is shown from the m/z range of 927 – 938 in order to visualize the presence or absence of the isotopic mass envelops for the GMP-labeled peptide and the corresponding ^15^N- and ^13^C-labeled GMP peptides (right panel). For the unlabeled sample, average mass spectrum for retention times of ∼45.68 – 45.78 is shown. The inset indicates the mass added to any peptide modified by GMP, ^15^N-GMP and ^13^C-GMP as well as the mass differences between the GMP-labeled and the heavy-labeled GMP peptides for the specified charge states (z). c,The top-scoring EThcD peptide spectrum match for same GMP-labeled peptide is shown with labeling of main fragment ions. Inset shows an abbreviated list of expected peptide mass fragments, which are colored red or blue if they were matched in the spectrum within an error of 0.04 Daltons. Note that not all matched ions are labeled in the spectrum.

For the next step of analysis, candidate peaks were subjected to MS/MS analysis in the tribrid mass spectrometer using higher energy collisional dissociation (HCD) fragmentation methodology as well as electron transfer/higher-energy collision dissociation (EThcD). EThcD often preserves the attachment of labile peptide modifications that are difficult to preserve with HCD alone ^15-17^. Table 1 summarizes all GMP-modified peptides identified by MS/MS using the Sequest HT database search algorithm (also see Supplementary Table 4) ^18,19^. In SARS-CoV-2, we could only verify a single GMP-modified site by MS/MS located at ser-2 of the nsp7 recombinant protein, which is the N-terminal ser-1 residue in the nsp7 protein produced by the virus. Multiple forms of this same modified peptide were identified. For EAV, five total sites were identified in nsp proteins. This included nucleotidyation of nsp7 lys-143, which was observed in multiple peptides, and one site in nsp9. All GMP linkages in the EAV nsp proteins occurred via a phosphoramidite (P-N) bond rather than the O-linked GMP found in SARS CoV-2 (Table 1).

**Table 1.**
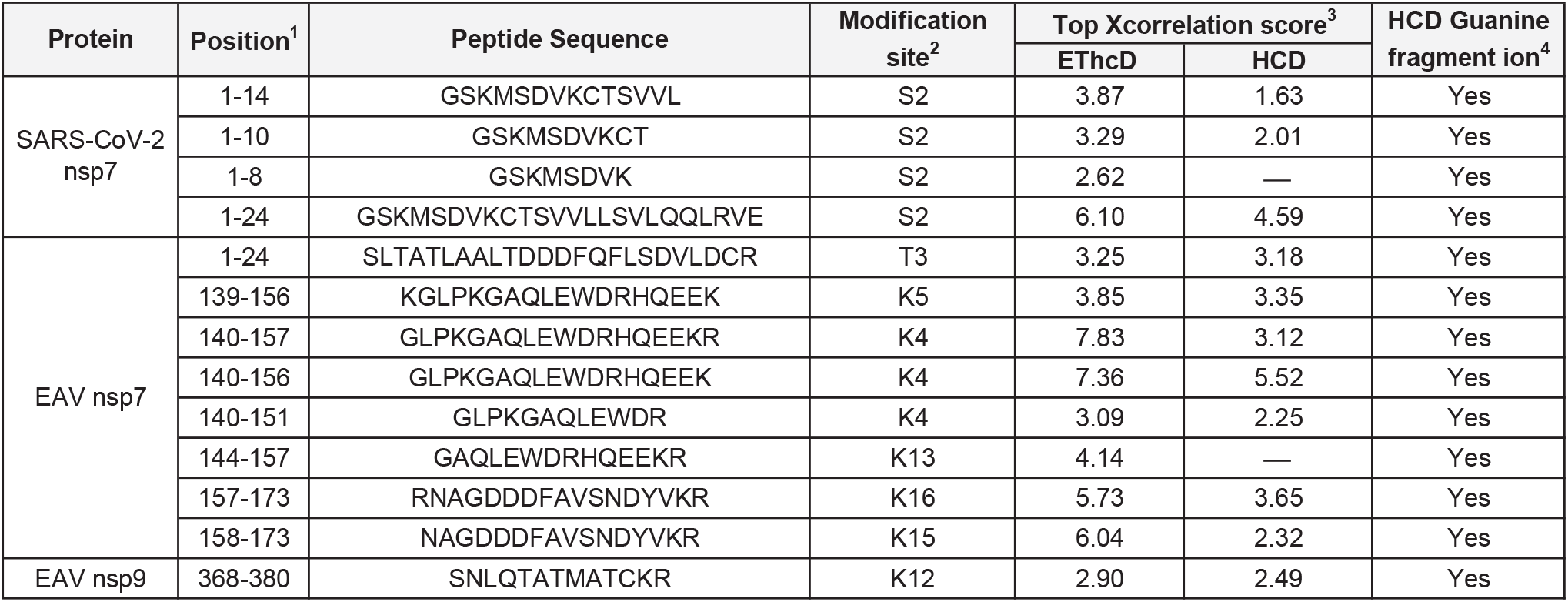
GMP-labeled peptides detected by LC-MS/MS after peptide spectrum matching with the Sequest HT database search algorithm. ^1^For SARS-CoV-2, positions are shown for the recombinant protein that contains an N-terminal glycine. The virally produced protein does not contain a glycine at position 1. ^2^Assigned GMP localization for the top-scoring spectrum match. ^3^Top score amongst GMP, ^15^N-GMP and ^13^C-GMP labeled peptides are shown. “-” indicates that corresponding HCD spectra did not pass MS/MS search criteria. ^4^Indicates guanine fragment ion was the predominant ion in corresponding HCD spectra, even when they were not identified as peptide spectrum matches using the MS/MS search criteria.

Compared to HCD, EThcD spectra provided clean, higher-quality, and more interpretable data that had improved spectrum matching scores (Xcorrelation scores) and that contained both N-terminal and C-terminal fragment ions (c-ion series and z-,y-ion series, respectively) (Fig. 2c, Fig. 3, Extended Data Fig. 6 - 9). Preservation of the GMP attachment allowed us to determine the modification site. However, some loss of the GMP modification during fragmentation was observed. One example is the 1438.73035 ion in Figure 2c that corresponds to the z^13+^ fragment that lost the GMP modification. If not for the presence of the y^13+^ ion at 1798.78857, it would be easy to assign the GMP attachment site to the N-terminal glycine residue instead.

**Figure 3.**
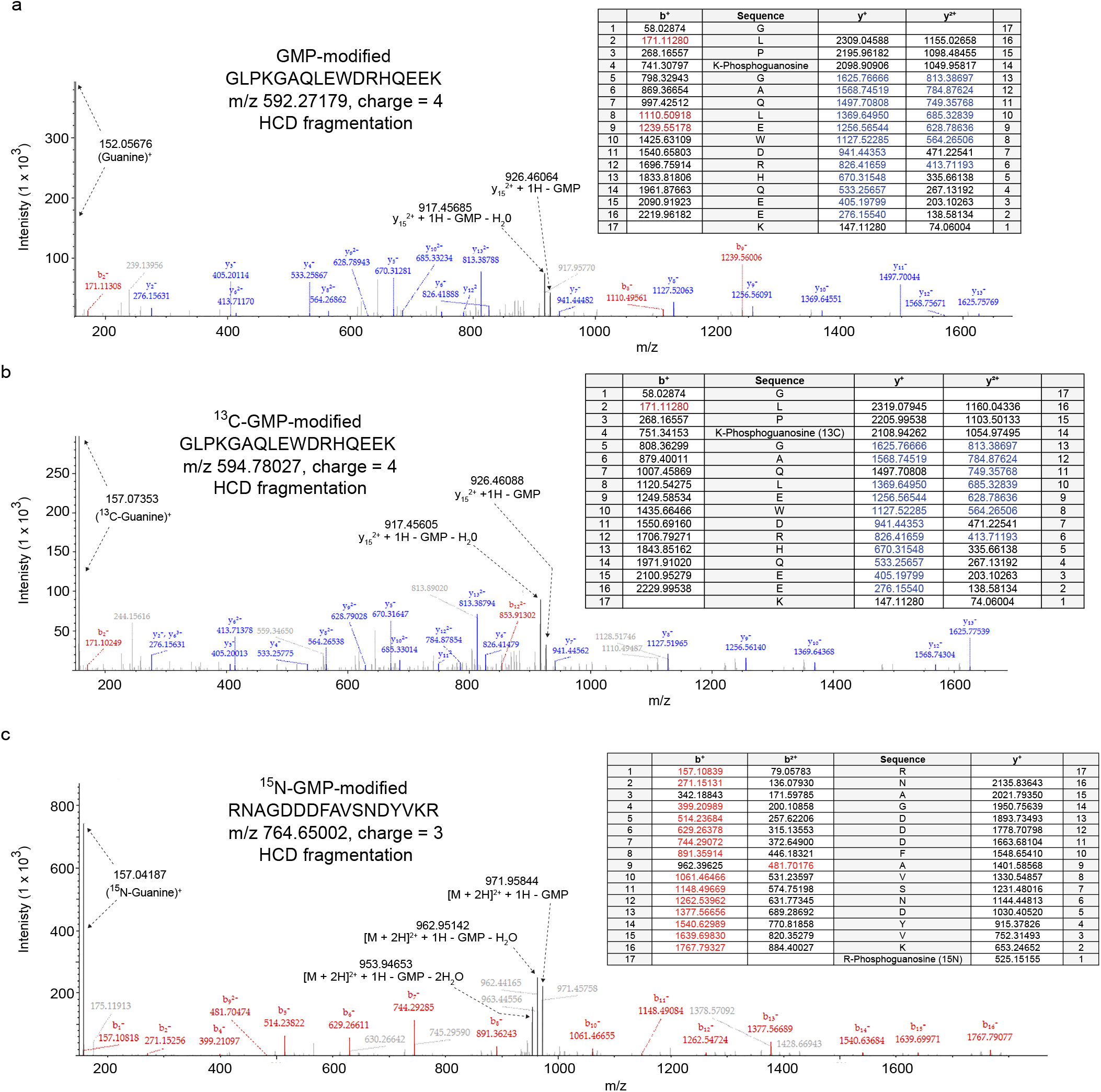
MS/MS spectra of GMP, ^15^N-GMP and ^13^C-GMP modified peptides fragmented with HCD. Examples of top-scoring HCD peptide spectrum matches from two EAV nsp7 peptides are shown that are either labeled closer to the N-terminus (a, b) or the C-terminus (c). Each spectrum represents a peptide modified with either GMP, ^15^N-GMP or ^13^C-GMP as indicated. Inset lists expected fragment ions in an identical manner as in Figure 2c. Spectra are additionally labeled with guanine fragmentions and dominant peptide fragments that have lost the GMP modified plus one or two H_2_O molecules. Note that Sequest HT assigned the GMP modification site for the spectrum in panel c to R17, instead of the correct site at K16.

The most prominent feature in HCD spectra was the fragmentation of the GMP moiety itself that, in every instance, leaves a characteristic and dominant guanine (C_5_N_5_OH_6_) tracer ion (Table 1, Fig. 3, and Extended Data Fig. 6-9). The guanine ions were also observed in EThcD spectra, but at a low intensity. The mass of the guanine fragment changed appropriately depending on whether the peptide was labeled with GMP, ^13^C-GMP or ^15^N-GMP. The theoretical masses of guanine that is derived from each of these versions of GMP are 152.0572, 157.0740 or 157.0424, respectively. Peptide ions that do not contain the modification were prominent in these HCD spectra. For example, when the modification was located closer to the N-terminus, the y-ion series is easily identified, and the b-ion series is missing (Fig. 3a, b). Similarly, when the modification is closer to the C-terminus, the b-ion series is prominent, and the y-ion series is generally absent (Fig. 3c). The missing ion series was often observed with the loss of the GMP modification (Extended Data Fig. 6 and Extended Data Fig. 10), which makes its localization difficult if HCD is used alone. One main ion that has lost the GMP modification along with one or two H_2_O molecules often stands outs, for example, the 926.46 and 917.45 m/z ions in Figure 3a and 3b, and the 971.96, 962.95 and 953.95 ions m/z in Figure 3c.

Until now, successful MS identification for GMP-peptide adducts has not been demonstrated. The results reported here allow a more systematic and high-throughput method to find such sites. We show that EThcD fragmentation is favorable over HCD fragmentation because both sets of the N- and C-terminal ion series are generated with the modification intact. Much of the energy provided in HCD fragmentation causes destruction of the GMP molecule, similar to what occurs with glycan PTMs in the collisional-induced dissociation (CID) fragmentation technique that is comparable to HCD ^20,21^. However, HCD can be a powerful diagnostic tool used to locate peptides that are GMP-modified because, without exception, the dominant MS/MS ion is always a guanine at a m/z of 152.0572. AMP-peptide adducts have been examined by MS/MS using CID ^22^. These spectra also featured destruction and loss of the nucleotide modification that could make peptide identification difficult. However, fragmentation of the AMP moiety left diagnostic ions at 136.1 and 250.1 that could be used to locate modified peptides in complex mixtures.

We failed to identify GMP-modified sites in SARS-CoV-2 nsp8, even though the proteins were clearly labeled with radioactivity in the gel-based assay. Although we utilized a variety of proteases to examine our samples by LC-MS, it is possible that the modified peptides could not easily be isolated under the liquid chromatography conditions or they failed to ionize efficiently, which are both necessary to produce a strong MS signal. GMP-modification added hydrophobicity to some peptides as they eluted with higher organic solvent on a reverse phase column compared to their unmodified counterparts (Fig. 2a vs 2b). It is also possible that the stability of GMP attachment is dependent on the chemical environment provided by neighboring amino acids. Additionally, attachment to amides through the formation of phosphoramide bonds is known to be unstable in acid conditions used in LC-MS (Extended Data Fig. 1) ^3,12,13^. Potentially, the observation of the GMP-modified ser-2 on recombinant SARS-CoV-2 nsp7 in the mass spectrometer could be the manifestation of a lys-3 GMP-modification via a phosphoramide bond that was transferred to a more energetically favorable position on the neighboring residue during the sample handling. It should be noted that non-canonical phosphorylation on basic amino acids, which uses identical phosphoramidate chemistry (i.e., the P-N bond versus P-O in S, T and Y amino acids) for the GMP linkage, is beginning to emerge as an important component of signaling systems in nature ^16,23^. These phosphoramidate bonds are more labile under the conditions commonly used in analytical instrumentation such as HPLC and mass spectrometry and are therefore understudied. This highlights the need for more comprehensive studies that examine the impact of additional factors on their in vivo and in vitro lability, such as the specific local chemical environment provided by unique amino acid sequences on either side of the modified residue.

Nucleotidyl transferase activity plays a role in diverse biological processes in bacteria, viruses and eukaryotes. The addition of AMP (AMPylation or adenylation) was first described as a mechanism to regulate *E. coli* glutamine synthetase activity ^24^. It is now recognized, along with UMPylation, as a post-translation modification that impacts bacterial DNA replication ^25,26^, ER and oxidative stress responses ^27-29^, bacterial pathogenesis ^30,31^, and viral RNA synthesis ^32^ where nucleotides are stably attached to serine, threonine or tyrosine resides ^33,34^. AMP and GMP adducts are also formed as intermediates in RNA capping ^12,35,36^, DNA ligation ^37,38^, SUMOylation and ubiquitinylation ^39-43^ as well as other enzymatic reactions ^44,45^. In these cases, as with non-canonical phosphorylation of lysine and histidine, the nucleotide forms less stable linkages with amine (lysine or histidine) or carboxyl groups that promote its transfer to substrate.

Here we have demonstrated nucleotidylation activity of SARS-CoV-2 proteins that is hypothesized to be critical for viral propagation ^3^. Mutation of nsp12 K73 caused a loss of transferase activity, which indicates the N-terminal NiRAN domain is necessary for formation of GMP adducts on nsp7 and nsp8. Importantly, the NiRAN domain (residues 1-250) is distant from the RNA polymerase active site (residues 400 – 932) and also the binding sites for nsp7 and nsp8 ^1,2^. Nsp7 and nsp8 also form alternate, oligomeric structures ^46^. Unlike other conserved nucleotidyl transferase domains, the NiRAN domain is newly defined and its functional role in viral life cycles remains to be elucidated. While many components of coronavirus RNA capping machinery have been characterized, the hypothesized guanyltransferase, which traditionally forms GMP-intermediates as observed here, remains undiscovered ^12,35,36,47^. In other RNA viruses, a uridylylated protein, known as VPg, is used to initiate RNA synthesis ^48-53^. SARS-CoV-2 nsp7 and nsp8 are positioned to interact with the RNA 5’ terminus when bound to nsp12 ^54,55^ and are critical for robust RNA replication ^9,56^. We expect the data provided here will lay the foundation for future studies defining the exact mechanistic and functional purposes of this novel nucleotidylation activity in SARS-CoV-2 and related coronaviruses. The translational utility of this information for discovering and/or designing new inhibitors of the process that function to thwart COVID-19 is an important future goal for research on SARS-CoV-2 as well as unknown but potentially newly emerging coronaviruses.

## Methods

### Expression and purification of the EAV and SARS-CoV-2 proteins

DNA for SARS-CoV-2 nsp12 encompassing the a.a. 1-932 and EAV nsp9 1-693 was synthesized with codon optimization (Genscript) and cloned into pFastBac with an N-terminal MG addition and C-terminal TEV protease site and two Strep tags. Expression was performed by transducing recombinant baculoviruses into Sf21 insect cells (Expression Systems). Cells were harvested by centrifugation and resuspended in 25 mM HEPES pH 7.4, 300 mM sodium chloride, 1 mM magnesium chloride, and 2 mM dithiothreitol. The resuspended cells were then lysed using a microfluidizer, clarified by centrifugation at 25,000 × g for 30 min and filtered using a 0.45 μm filter. Nsp12 was purified using Streptactin Agarose (IBA) eluting with 2 mM desthiobiotin. Eluted protein was further purified by size exclusion chromatography using a Superdex200 column (GE Life Sciences) in 25 mM HEPES pH 7.4, 300 mM NaCl, 0.1 mM magnesium chloride, and 2 mM tris(2-carboxyethyl)phosphine. Full-length, codon-optimized nsp7 and nsp8 genes were cloned into pET46 for expression in E. coli. The N-terminal tags for SARS-CoV-2 nsp7 and nsp8 are MAHHHHHHVDDDDKMENLYFQG and for EAV nsp7 is MAHHHHHHVGTENLYFQ. The TEV protease cleavage sites (ENLYFQ|G) were positioned to leave N-terminal glycines on SARS-CoV-2 nsp7 and nsp8 and no additional amino acids on EAV nsp7. Proteins were expressed in Rosetta2 pLysS E. coli (Novagen). Bacterial cultures were grown to an OD_600_ of 0.8 at 37 °C, and then the expression was induced with a final concentration of 0.5 mM of isopropylβ-D-1-thiogalactopyranoside and the growth temperature was reduced to16 °C for 14–16 h. Cells were harvested by centrifugation and were resuspended in 10 mM HEPES pH 7.4, 300 mM sodium chloride, 30 mM imidazole, and 2 mM dithiothreitol. Resuspended cells were lysed using a microfluidizer. Lysates were cleared by centrifugation at 25,000 × g for 30 min and then filtered using a 0.45 μm filter. Protein was purified using Ni-NTA agarose (Qiagen) eluting with 300 mM imidazole. Eluted proteins were digested with 1% (w/w) TEV protease. TEV protease-digested proteins were passed over Ni-NTA agarose to remove uncleaved proteins and then further purified by size exclusion chromatography using a Superdex200 column (GE Life Sciences) in 25 mM HEPES pH 7.4, 300 mM sodium chloride, 0.1 mM magnesium chloride, and 2 mM dithiothreitol.

### Nucleotidylation assay

Nucleotidylation assays were performed similarly to as previously described ^1^. The EAV nsp9 or SARS-CoV-2 nsp12 RdRP proteins were incubated with a 2 – 4 molar excess of EAV nsp7 or SARS-CoV-2 nsp7 and nsp8 proteins, respectively, in following buffer: 50 mM Tris pH 8.5, 6 mM MnCl_2_, 1 mM DTT, 25 mM NaCl and 2% glycerol. Labeling reactions were initiated by the addition of GTP and were incubated for 30 minutes at 30°C. Radioactivity assays employed 0.5 – 5 μCi α-^32^P-GTP (Perkin Elmer) with total GTP concentration of 0.17 μM, whereas mass spectrometry assays used 0.2 – 1 mM GTP, ^15^N-GTP (Sigma), or ^13^C-GTP (Sigma).

### SDS-PAGE and autoradiography

Samples were mixed with 4x LDS loading dye (Thermo) containing 200 mM DTT and analyzed on Bolt Bis-Tris Plus gels (Thermo) according to manufacturer’s instructions. Gels were either stained with GelCode^™^ Blue Protein Stain (Fisher Scientific) or prepared for autoradiography by incubation in a 10% polyethyleneglycol 8000 (Sigma) solution for 30 minutes and gel drying at 70°C for 45 minutes using a Hoefer gel drier, Savant condenser unit and an TRIVAC pump (Leybolt). Gels were exposed to phosphor screens for 2 to 48 hours and developed with a Molecular Dynamics Storm 850 imager. Gel images were examined using ImageJ opensource software.

### LC-MS/MS

Nucleotidylation reactions were brought to 6.7 M Urea in 50 mM ammonium bicarbonate, 5 mM DTT and incubated at 42°C for 15 minutes. Cysteines were alkylated by the addition of 15 mM iodoacetamide for 30 minutes at room temperature. Following the neutralization of unreacted iodoacetamide with 15 mM DTT, nsp proteins were diluted to a final concentration if 1M urea and digested overnight with either Trypsin/LysC mix, Chymotrypsin, or GluC (Promega) according to manufacturer’s instructions. The resulting peptides were desalted using OMIX C18 pipette tips (Agilent Technologies) or 1 mL C18 Sep Pak cartridges (Waters) in 10 mM ammonium formate at pH 7.0, eluted in 75% acetonitrile, and dried to completion with a vacuum centrifuge.

For LC-MS/MS analysis, each digested sample was suspended in 0.1% formic acid and maintained at 7°C until analysis within 12 hours. Samples were loaded onto Thermo Scientific Easyspray C4 or C18 nanocolumn and eluted with a gradient to 80% acetonitrile / 0.1% formic acid at 300 nL/min at room temperature using an Ultimate 3000 series liquid chromatography system. Eluted peptides were ionized in-line with a Thermo Scientific Easy Spray source for direct analysis with a Thermo Scientific Fusion Lumos Orbitrap mass spectrometer and subjected to a targeted or data-dependent MS/MS acquisition scheme that collected both HCD and EThcD spectra in the high resolution orbitrap.

LC-MS peaks were defined by analyzing the data with the feature mapping and precursor quantification nodes in Proteome Discoverer 2.4 that determined the retention time, charge state, and abundance of every peak in each searched file. The data were filtered and exported to Microsoft Excel to search for peaks that 1) were not present in unlabeled samples and 2) that possessed two peaks with the same charge state at a similar retention time (+/- 0.5 minute) and were mass-shifted by 4.9852 or 10.0336 amu [MH+ ion masses] for ^15^N- or ^13^C-GMP labeling, respectively, within a +/- 1.5 ppm error limit (Extended Data Figure 4, Extended Data Figure 5). Supplementary Table 1, Supplementary Table 2, and Supplementary Table 3 are the Microsoft Excel worksheets used to perform these searches once the data was exported.

To identify peptide spectrum matches, data were searched with the Sequest HT algorithm through Proteome Discoverer 2.4 and matched with a mass error tolerance of 0.04 Da for MS/MS fragment ions ^2^. GMP adducts were assigned by allowing a dynamic modification for the mass addition of 345.0474 (and 350.0326 or 355.0810 for ^15^N and ^13^C labeled GMP) to H, S, T, Y, K or R, based on known phosphodiester or phosphoramide attachment chemistry ^3-5^, although all possible sites were considered in initial, preliminary searches. For the SARS-CoV-2 GMP localization on nsp7, initial searches assigned the modification site to the N-terminal glycine at peptide position 1, but the modification site was manually determined to be located at ser-2 (ser-1 in natural nsp7) because of the presence of the y13+ ion (ser-2) at an m/z of 1798.78857 that contained the GMP moiety. For each unique peptide m/z, the top-scoring peptide spectrum matches for both EThcD and HCD are presented in Supplementary Table 4 along with GMP-attachment site assignments and additional information. The top-scoring, overall match for each peptide is presented in Table 1 along with the corresponding attachment site assignment. All matches had a precursor mass error of +/- 1.5 ppm or less and were validated with a target-decoy search using a cut-off false discover rate value of 1%. Only peptides with a top spectrum match Xcorrelation score of 2.5 or greater were reported. Extracted ion chromatograms and spectra were generated using Freestyle software (Thermo Scientific). Note that for EAV reactions, not all GMP-modified candidate peaks were examined in MS2, because all EAV data was acquired prior to data analysis and MS2 was performed in a data-dependent acquisition scheme. Though a single site was found in SARS-CoV-2, additional phosphoramidate linked GMP sites likely exist that may not have been observed because of the lability of the GMP attachment in the acidic conditions used for LC-MS/MS (Extended Data Figure 1).

### Data availability

The raw mass spectrometry datasets and proteome discoverer results that were generated and analyzed in the current study are available through the MassIVE repository under the identifier MSV000085857 [https://doi.org/doi:10.25345/C52B2B]. All figures were derived from this data except for Figure1, Extended Data Figure1, and Extended Data Figure 2.

## Acknowledgments

Funding to support this work was provided to R.N.K. from NIH/NIAID (Grant No. AI123498) and M.R.S. from NSF (Grant No. MCB-1713899).

## Author contributions

B.J.C designed and performed the experiments and wrote the manuscript. R.N.K. purified and provided the proteins, conceived the project, helped to design experiments, and contributed to the writing of the manuscript. M.R.S. conceived the project, helped to design experiments, and wrote the manuscript.

## Additional information

Supplementary Information is available for this paper.

**Extended Data Figure 1.**
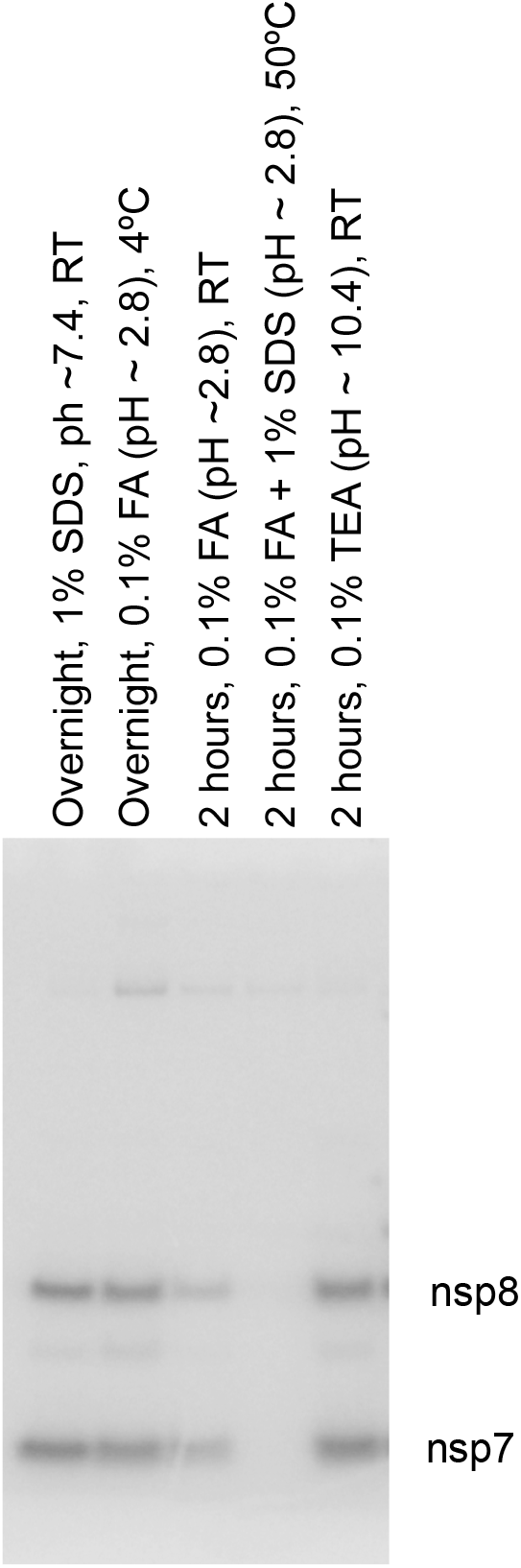
Stability of GMP covalent attachment on SARS-CoV-2 nsp7 and nsp8. SARS-CoV-2 proteins nsp7 and nsp8, radiolabeled with ^32^P-GTP, were incubated in the indicated conditions and were analyzed by SDS-PAGE. Autoradiography was used to visualize radioactive proteins. SDS was necessary in the heated sample to prevent aggregation. Abbreviations: FA, formic acid; TEA, triethylamine; RT, room temperature.

**Extended Data Figure 2.**
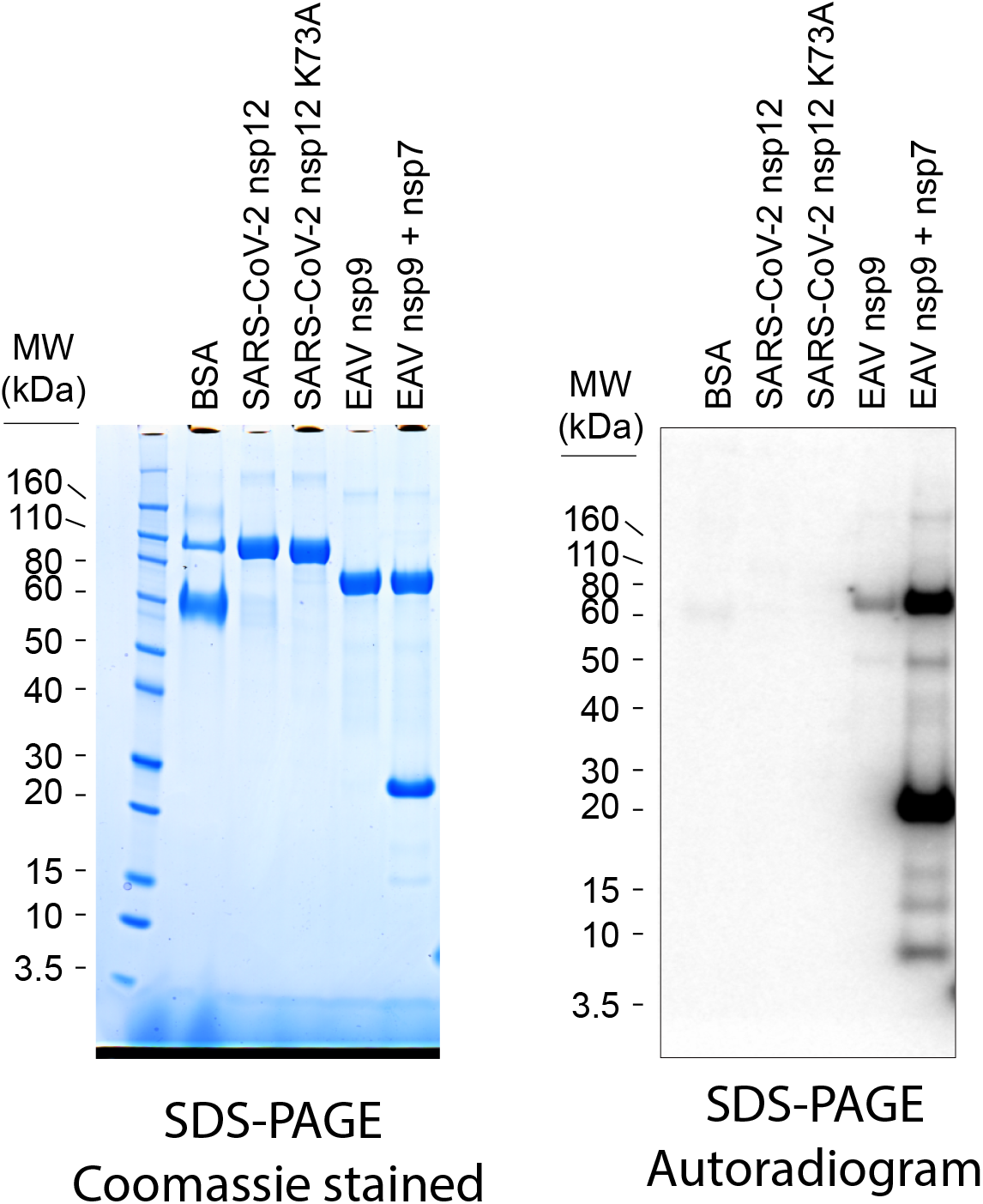
Presence of EAV nsp7 increases nsp9 nucleotidylation. The indicated proteins were subjected to a nucleotidylation reaction and analyzed by SDS-PAGE followed by Coomassie staining to visualize total proteins (left panel) or autoradiography to visualize radiolabeled proteins (right panel).

**Extended Data Figure 3.**
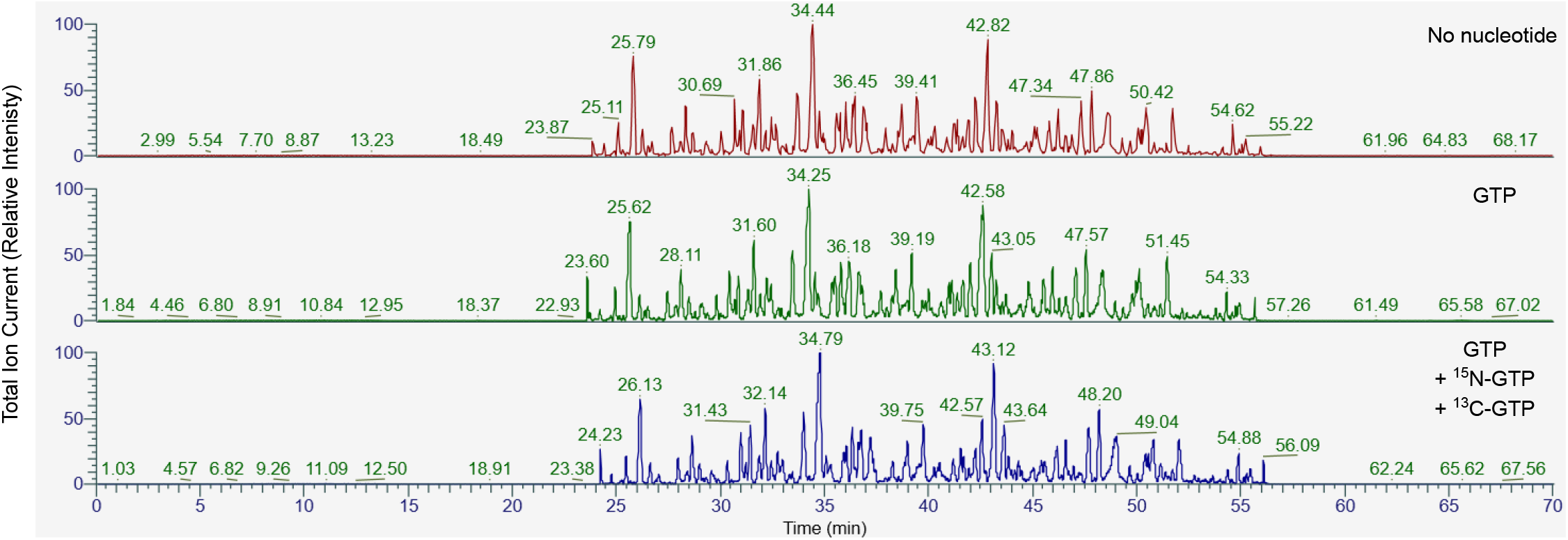
Total Ion Chromatrograms for SARS-CoV-2 peptide samples. The total ion chromatrograms for Figure 2a and 2b are shown that were subjected to nucleotidylation reactions, as described in the text, and digested with chymotrypsin. The overall peak profiles were similar and peptides eluted at similar retention times between 24 and 57 minutes.

**Extended Data Figure 4.**
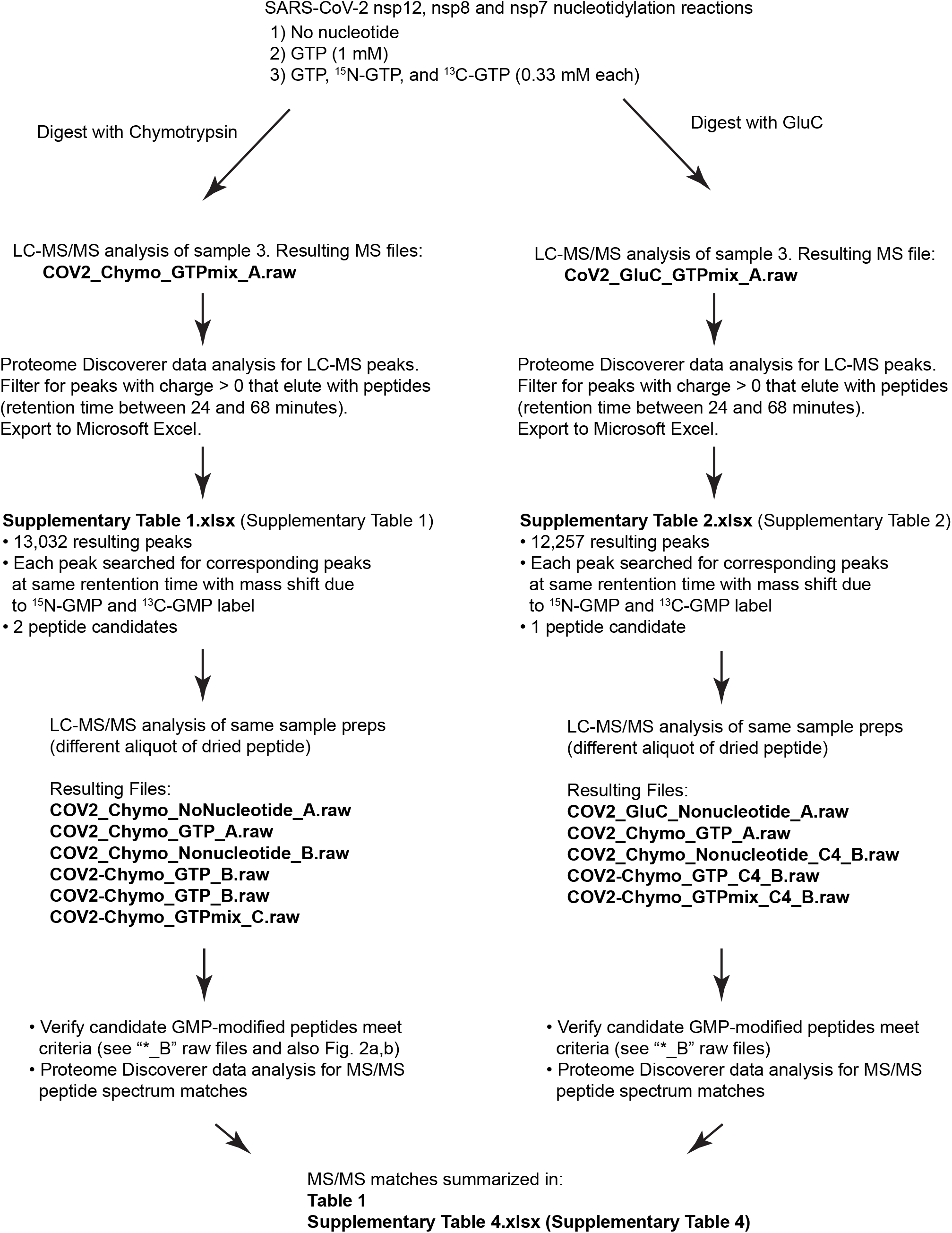
Workflow for identification of SARS-CoV-2 GMP-labeled peptides. See indicated files and tables for raw data, analysis and results.

**Extended Data Figure 5.**
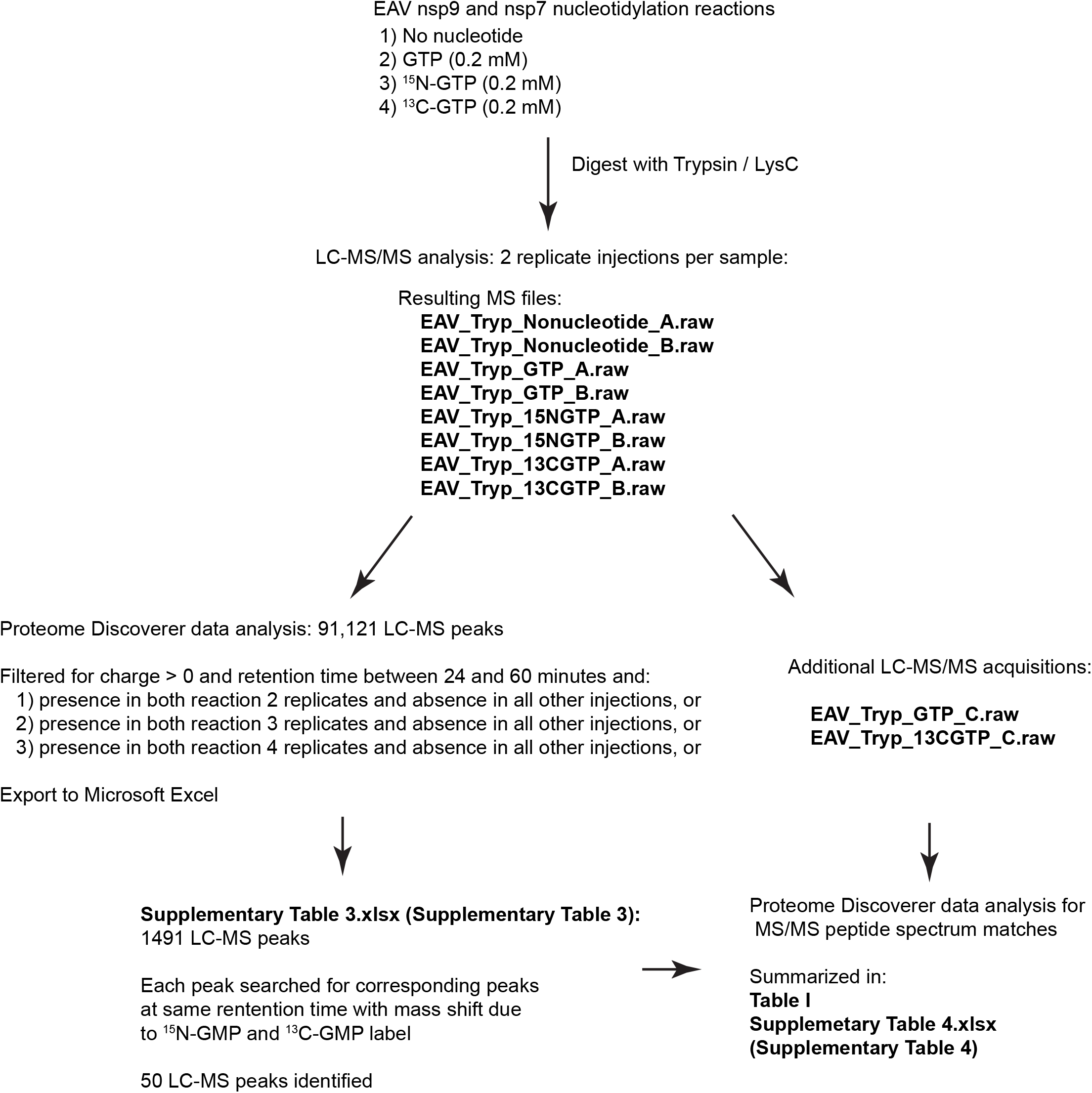
Workflow for identification of EAV GMP-labeled peptides. See indicated files and tables for raw data, analysis, and results.

**Extended Data Figure 6.**
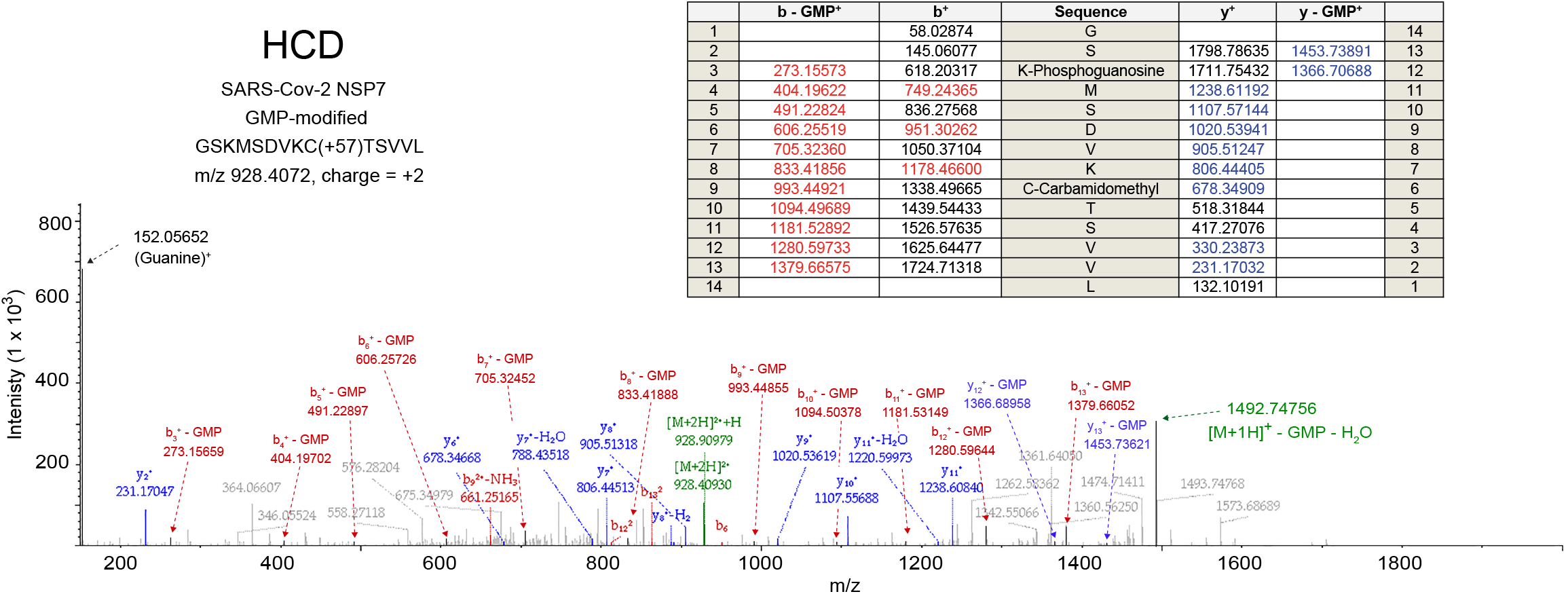
HCD spectrum for GMP-modified SARS-CoV-2 nsp7 peptide 1-14. The top scoring HCD spectrum match for GMP-modified nsp7 peptide is shown for direct comparison to EThcD spectrum in Fig. 2c. Note that the modification site was assigned to K3 by Sequest HT for this spectrum, instead of S2. Guanine and fragments that lost the GMP modification were manually labeled after inspection.

**Extended Data Figure 7.**
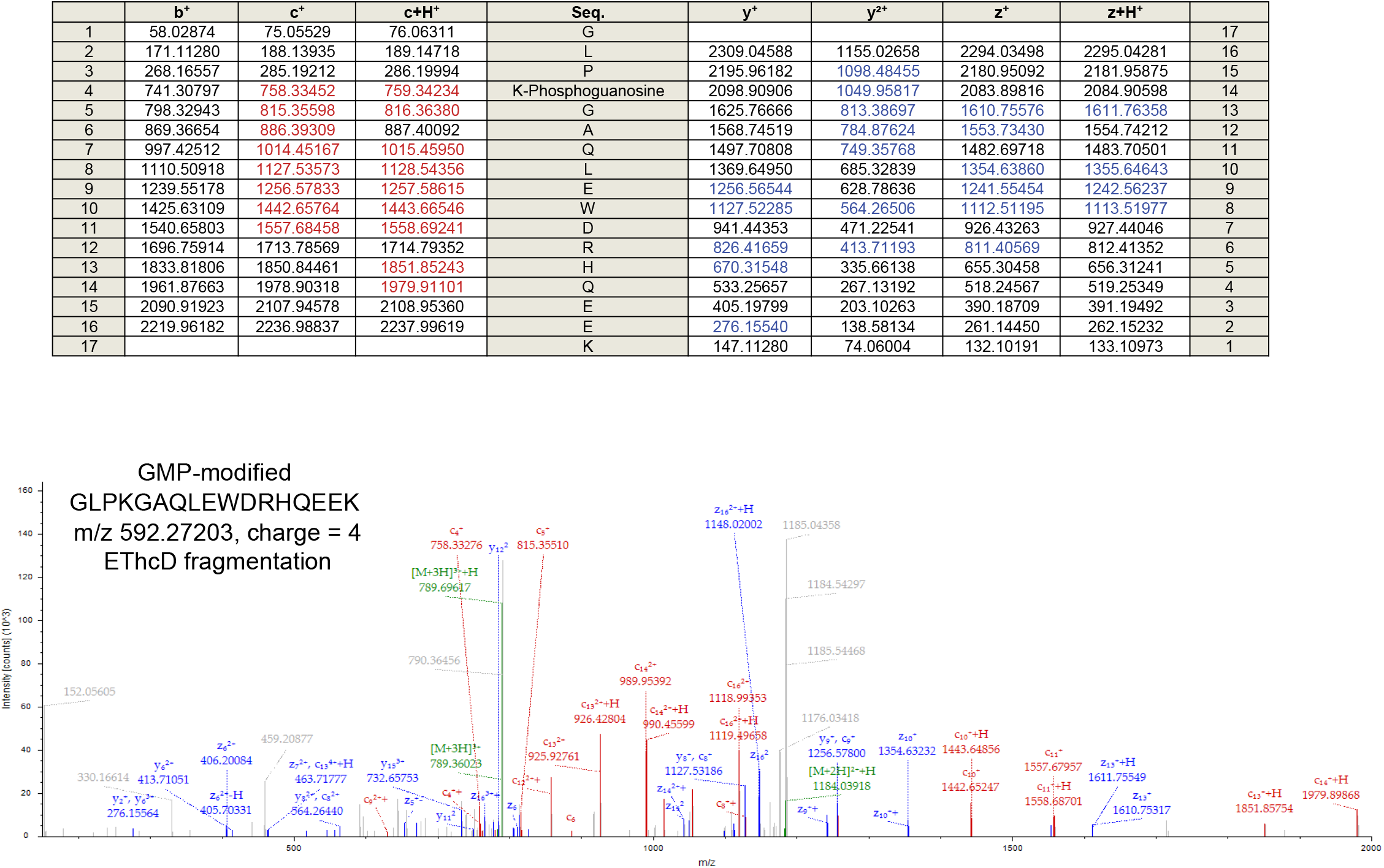
EThcD spectrum for GMP-modified EAV nsp7 peptide 140-156. The top scoring EThcD spectrum match for the GMP-modified nsp7 peptide in Figure 3a is shown.

**Extended Data Figure 8.**
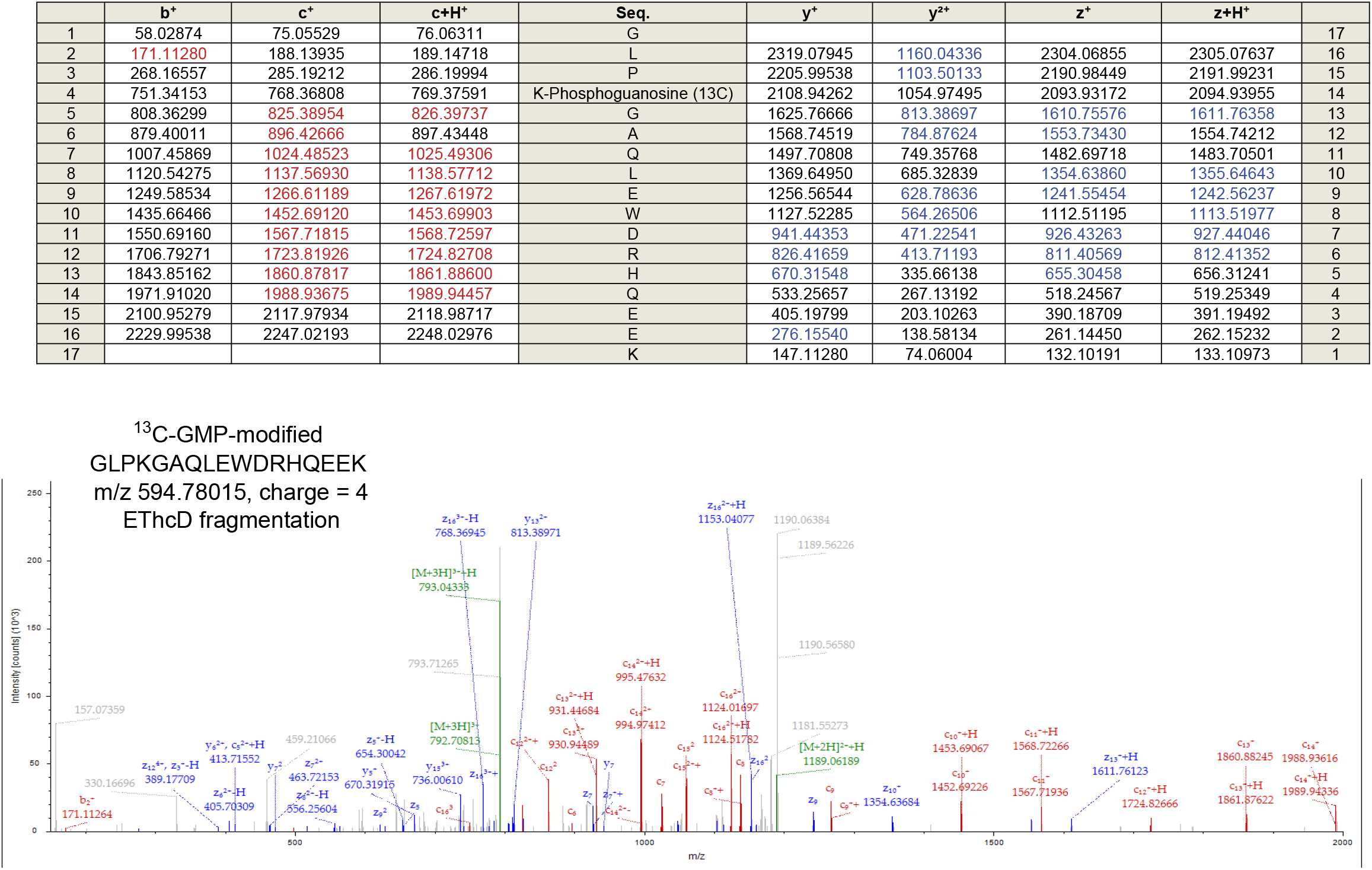
EThcD spectrum for ^13^C-GMP-modified EAV nsp7 peptide 140-156. The top scoring EThcD spectrum match for the ^13^C-GMP-modified nsp7 peptide in Figure 3b is shown.

**Extended Data Figure 9.**
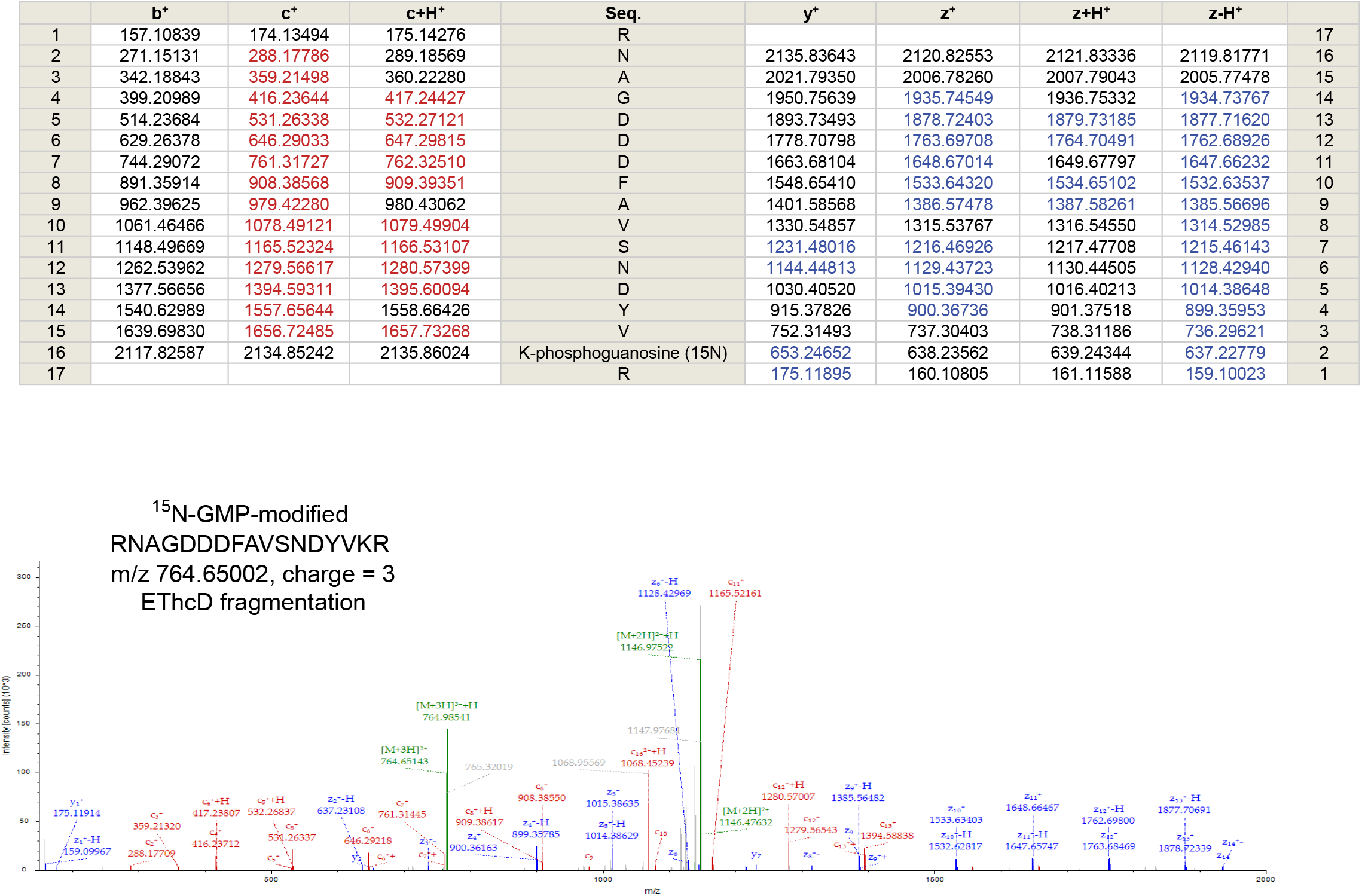
EThcD spectrum for ^15^N-GMP-modified EAV nsp7 peptide 157-173. The top scoring EThcD spectrum match for the ^15^N-GMP-modified nsp7 peptide in Figure 3c is shown.

**Extended Data Figure 10.**
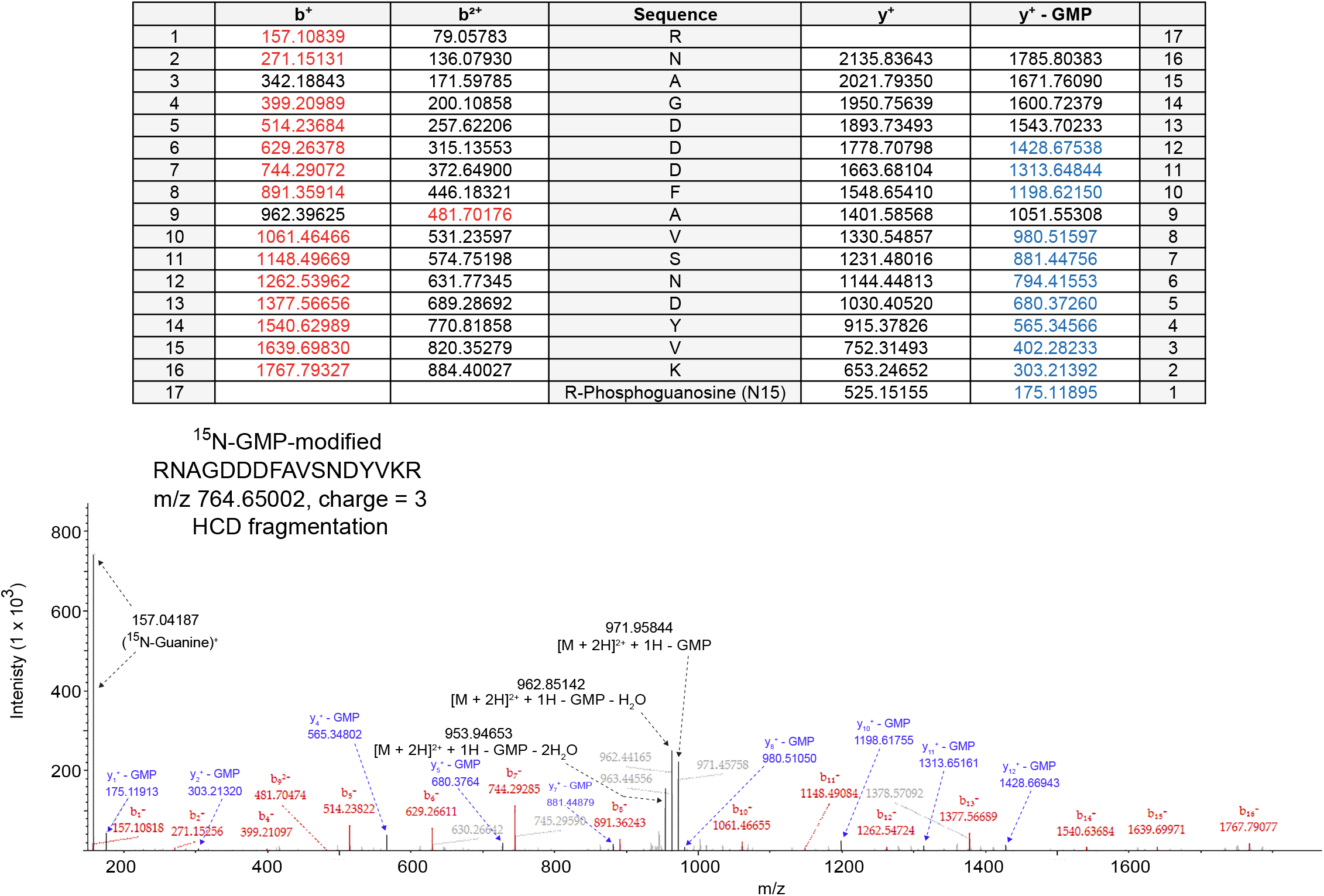
HCD spectrum for ^15^N-GMP-modified EAV nsp7 peptide 157-173. The top scoring HCD spectrum match for the ^15^N-GMP-modified nsp7 peptide in Figure 3c is relabeled to include fragment ions that have lost the GMP modification.

## Supplementary Information Guide

**Supplementary Table 1:** Data analysis of LC-MS peaks for a single SARS-CoV-2 chymotrypsin-digested sample that contained peptides labeled with GMP, ^15^N-GMP and ^13^C-GMP. “Search set up” tab shows formulas and criteria to search each LC-MS peak against all others (“index to search” tab). Two additional tabs show search results and filtered results.

**Supplementary Table 2:** Data analysis of LC-MS peaks for a single SARS-CoV-2 GluC-digested sample that contained peptides labeled with GMP, ^15^N-GMP and ^13^C-GMP. “Search set up” tab shows formulas and criteria to search each LC-MS peak against all others (“index to search” tab). Two additional tabs show search results and filtered results.

**Supplementary Table 3:** Data analysis of LC-MS peaks for EAV trypsin-digested samples that were not labeled with GMP or were labeled with either GMP, ^15^N-GMP or ^13^C-GMP. Peaks unique to two labeled, replicate injections (GMP-, ^15^N-GMP or ^13^C-GMP-labeled) were exported and searched against all others (“index to search” tab). Three tabs total.

**Supplementary Table 4:** Summary of all MS/MS peptide spectrum matches containing a GMP, ^15^N-GMP, or ^13^C-GMP adduct. Top scoring hits for each peptide mass (m/z) and for each fragmentation method (HCD or EThcD) are shown along with peptide sequence, Xcorrelation score, precursor mass error, raw data file name, scan number, and other relevant information.

